# Late B lymphocyte action in dysfunctional tissue repair following kidney injury and transplantation

**DOI:** 10.1101/343012

**Authors:** Pietro E Cippà, Jing Liu, Bo Sun, Sanjeev Kumar, Maarten Naesens, Andrew P McMahon

## Abstract

The mechanisms initiating the late immune response to allografts are poorly understood. Through transcriptome analysis of serial protocol biopsies in kidney transplant recipients, we found a tight correlation between the initial response to kidney injury and a late B lymphocyte signature associated with renal dysfunction and fibrosis, suggesting a link between dysfunctional repair and immunoreactivity. To specifically investigate the immunological consequences of dysfunctional repair, we followed the mouse kidney up to 18 months after ischemia/reperfusion. Even in the absence of foreign antigens we identified a sustained immune response in conjunction with the transition to chronic kidney damage. This tissue-driven immunological process involved both the innate and the adaptive immune system and eventually induced an antigen-driven proliferation, selection and maturation of B lymphocytes into broadly-reacting antibody secreting cells. These findings reveal an unappreciated role of dysfunctional tissue repair on local immunoregulation with a particular relevance for late transplantation immunobiology.

## Introduction

The immune system participates in tissue repair with contrasting effects: inflammation after injury is important to initiate the repair response but immune cells can contribute to secondary tissue damage.^1^^-^^5^ Organ transplantation is an interesting model to investigate the interaction between tissue injury and local immune regulation. Ischemia/reperfusion injury inevitably occurs during organ transplantation and triggers the coordinated activation of the innate and the adaptive immune system of the host in a complex immunological process leading to acute allograft rejection.^6^^-^^8^ This process has been extensively investigated in experimental models and can be effectively prevented and treated with currently available immunosuppressive drugs.^9^

The early and late immune responses to allografts are distinct processes: chronic forms of rejection remain mechanistically poorly understood and are not effectively treatable.^10^^-^^12^ As a result, long-term outcome after kidney transplantation has not substantially improved over two decades: approximately 4-5% of renal grafts are lost annually beyond the first year after transplantation, mainly because of late forms of immune-mediated injury (*chronic rejection*).^12^^-^^14^ Improved pathologic and immunologic diagnostic tools (e.g. C4d stain and detection of anti-HLA antibodies) indicate a critical role for B lymphocytes and donor specific antibodies in the late immune response to allografts. ^15,16^ However, at a time when the criteria for chronic antibody-mediated rejection are met, even aggressive immunosuppressive therapy does not substantially improve graft survival.^10,17^ The mechanisms initiating a donor-specific immune response, several months/years after transplantation and often without any clear precipitating event, are unclear and clinically important. In this study, we took advantage of recent technical advances in the characterization of the adaptive immune system and molecular processes determining the transition from acute to chronic kidney injury to investigate the impact of dysfunctional kidney repair on the late immune response following kidney injury and kidney transplantation.

## Results

### Chronic kidney injury and late B lymphocyte activity correlate after kidney transplantation

Transcriptional profiling of protocol biopsies from 42 kidney allografts in the first year after transplantation enabled an examination of changing gene activity in the post-transplant kidney. We identified a cluster of strongly correlated genes associated with fibrosis (e.g. COL1A1, DPT, MMP7), immunity (e.g. CD52, CXCL10, CCL21) and B cell action (immunoglobulin genes) (**Fig. 1a**). In agreement with previous reports, B cell associated transcripts and immunoglobulin genes increased over time after transplantation (**Fig. 1b-d**).^18,19^ The sample cluster analysis based on B cell-associated genes expressed in tissue biopsies collected at 12 months following transplantation indicated that the B cell signature was restricted to a subset of patients (**Fig. 1e**). Independently, t-distributed stochastic neighbor embedding (t-SNE) analysis identified a group of patients separating from the remaining study population by virtue of the higher expression of genes associated with fibrosis, wound healing and immunity (**Fig. 1f-h, Suppl. Table 1**). Since fibrosis and chronic inflammation are universal features of the maladaptive repair processes leading to chronic organ damage,^1,20,21^ we defined this group of patients as a maladaptive injury repair group (MIR-group). The MIR-group substantially overlapped with the subset of patients classified in the cluster analysis based on B cell-associated genes (odds ratio for the overlap 45.5, P<0.0001, **Fig. 1e**). Thus, the fibrosis and B cell signature show a strong concordance in kidney tissue 12 months after transplantation. These findings suggest two non-exclusive scenarios: (1) the host immune response to the allograft causes kidney damage, (2) maladaptive kidney repair triggers a late B cell response targeting the injured tissue.

**Figure 1.**
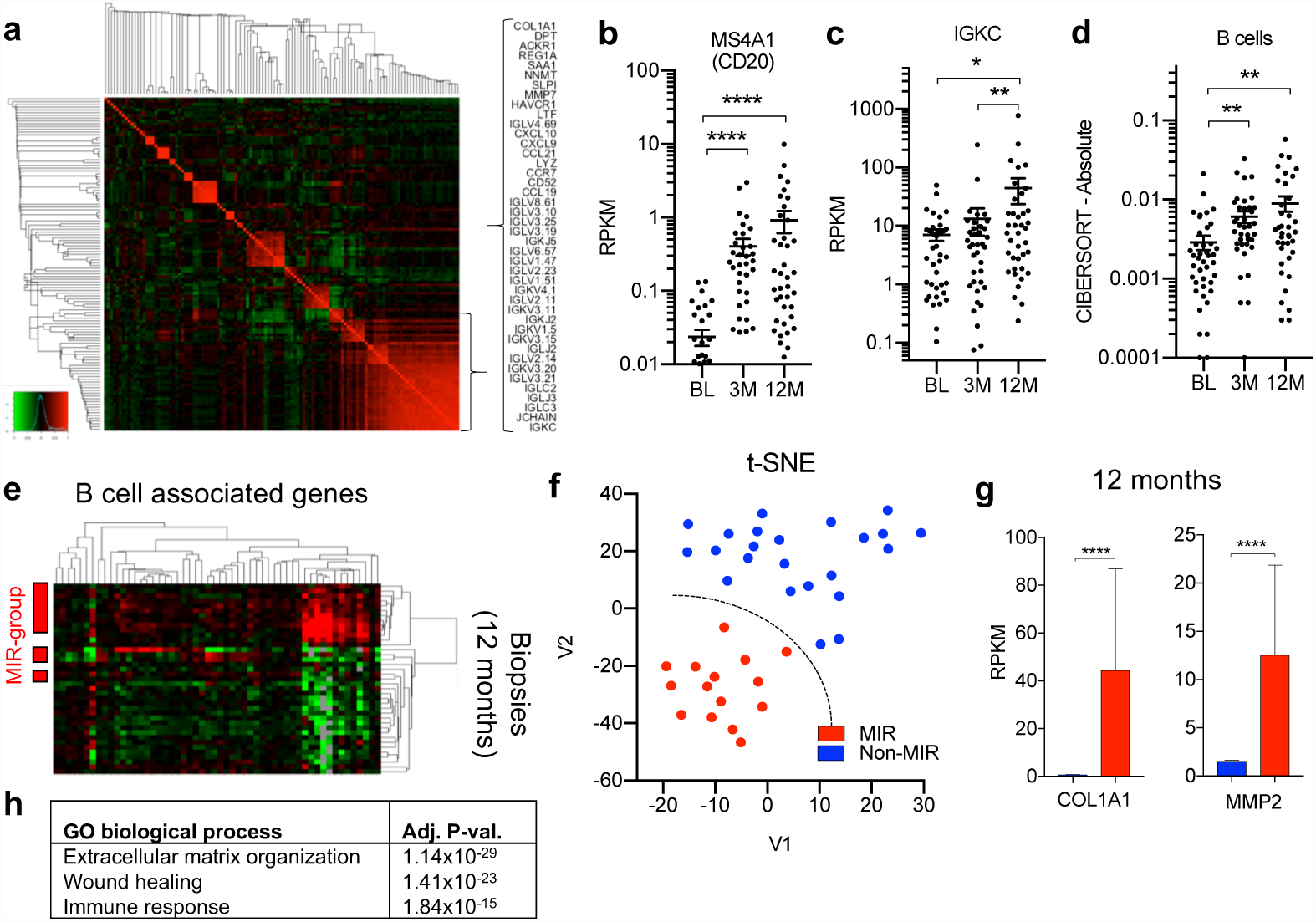
Chronic kidney injury and B cell immunity in human allografts. (a) Heatmap showing the gene expression correlation of the 120 most variably expressed genes in kidney biopsies collected at 3 and 12 months after transplantation. The names of the genes included in the cluster in the bottom right corner are shown. (b-c) RPKM values of MS4A1 (CD20) and IGKC over time in human kidney allograft biopsies. BL: baseline. N=39-42 for each time point. **** P<0.0001, **P<0.01, *P<0.05. (d) Semi-quantitative evaluation of immune cell infiltrates in the kidney at different time points after transplantation as determined by CIBERSORT analysis on RNAseq data. N=39-42 for each time point, **P<0.01. (e) Cluster analysis based on the expression of B cell associated genes including kidney biopsies collected at 12 months after transplantation.Patients classified in the MIR group are indicated on the left. (f) t-SNE analysis on RNAseq data from kidney biopsies collected 12 months after transplantation defining the classification of patients in the MIR group. (g) RPKM values of COL1A1 and MMP2 shown as examples of genes differentially expressed in MIR and non-MIR. ****P<0.0001. (h)Gene enrichment analysis including genes differentially expressed in the MIR group compared to non-MIR.

### Kidney injury precedes B cell mediated immunity in human allografts

To find evidence to distinguish between these alternative possibilities, we examined earlier biopsies from the patient cohort to examine the transcriptional response in the same kidney over time for factors that might precede the development of chronic kidney injury and late B cell activity signatures. We compared the MIR-group (as defined at 12 months after transplantation, **Fig. 1f**) with the rest of the study population referred to as the “non-MIR” group. At baseline, clinical characteristics of the patients and transcriptional profiles of MIR and non-MIR kidney biopsies were indistinguishable (**Suppl. Table 2**). Three months post-transplant, renal function was similar between the two groups. No histological evidence was found for lymphocytic infiltrates compatible with rejection (similar t, i and v scores according to Banff classification)^22^ or chronic tissue damage (e.g. fibrosis as determined by the ci score) in the MIR group (**Fig. 2a-b**). However, MIR group patients expressed higher levels of well-characterized markers of acute kidney injury and repair (e.g. LCN2, SOX9, ALDH2A1; **Fig. 2c**),^23,24^ suggesting a more pronounced or sustained response to tissue injury.

**Figure 2.**
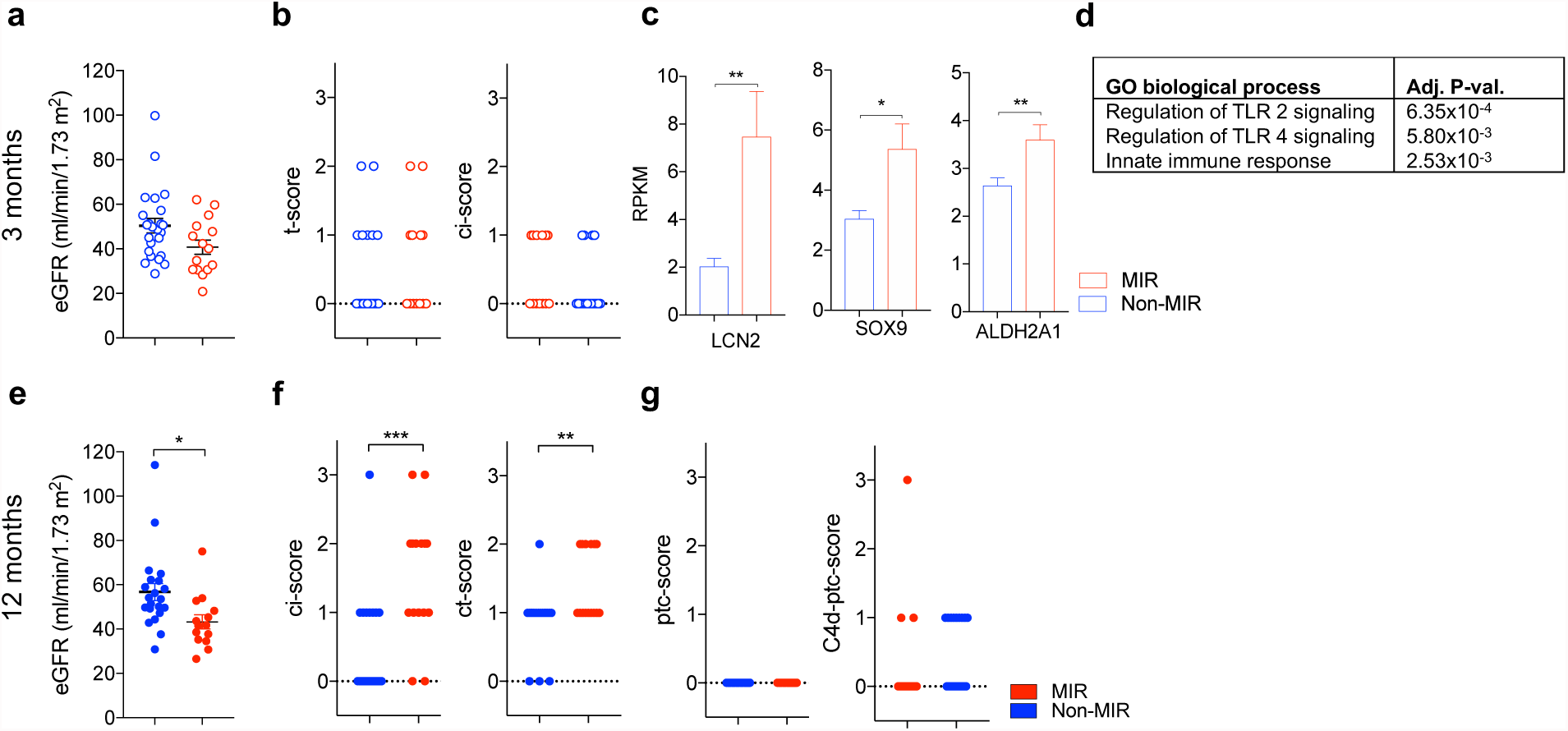
B cell action following tissue injury in human allografts. (a,e) Scatter plot showing the estimated glomerular filtration rate (eGFR) at 3 months (a) and 12 months (e) in kidney transplant recipients classified in the MIR and non-MIR group at 12 months. *P<0.05. (b,f,g) Scatter plot showing the histological evaluation of protocol biopsies according to Banff at 3 months (b) and 12 months (f,g). t: acute tubular lesions, ci: chronic interstitial lesions, ct: chronic tubular lesions, ptc: peritubular capillaritis, C4d-ptc: Cd4 positivity in peritubular capillaries. **P<0.01. (c) Histograms showing RPKM values of representative genes related to kidney injury and repair from kidney biopsies collected at 3 months after transplantation. *P<0.05, **P<0.01. Red symbols indicate patients of the MIR-group, blue symbols of the non-MIR group.Empty and filled symbols show data acquired at 3 and 12 months after transplantation respectively. (d)Gene enrichment analysis including genes with a higher expression in the MIR group at 3 months, among genes previously associated with acute rejection episodes.

To find evidence for subclinical rejection episodes as a potential cause of tissue injury in the MIR-group, we used an extensive list of genes associated with acute rejection that have been reported to detect rejection with greater sensitivity than conventional histology.^25^ Among the 186 analyzed genes, only 10 were detectably up-regulated in the MIR group (**Suppl. Table 3**). These genes relate to innate immunity (**Fig. 2d**), whereas genes more specifically linked to rejection and adaptive immunity were similarly expressed in the two groups. In contrast, among a set of 29 genes previously associated with acute kidney injury in kidney transplant recipients,^26^ 19 (65%) were significantly up-regulated in the MIR group and only 1 in the rest of the study population (**Suppl. Table 4**). Thus, kidneys exhibiting a signature of fibrosis and B cell activity at 12 months showed evidence for kidney injury, but not allograft rejection at 3 months.

One year after transplantation, patients in the MIR-group displayed a lower renal function and higher scores related to chronic kidney damage, consistent with transcriptional profiling (ci and ct score, **Figs. 2e-f**). In contrast, no histological evidence was observed for B cell mediated immunity as a cause of chronic kidney damage (absence of peritubular capillaritis and C4d staining, **Fig. 2g**).^27^ Histological analysis at 12 months revealed no evidence of increased inflammation in the MIR-group, pointing to a higher sensitivity for transcriptome analysis. In summary, a transcriptional signature of acute kidney injury at 3 months was associated with chronic organ damage and dysfunction at 12 months. The clinical data suggested that fibrosis in MIR patients did not result from a pathological immune response to the antigenically distinct allograft but support an alternative hypothesis in which the late immune response reflects a dysfunctional immune response related to ongoing tissue injury.

### Ectopic lymphoid tissue following renal ischemia-reperfusion injury in mice

To explore this hypothesis beyond the descriptive association provided by the clinical data, we turned to a mouse model where ischemia/reperfusion injury (IRI) led to a dysfunctional repair process that transitioned to chronic kidney disease over months, to specifically characterize the long-term immunological sequelae of maladaptive kidney repair.^28^ Sixteen months after a single IRI, the tissue architecture of the kidney was markedly distorted with flattened epithelial cells, cyst formation, extensive fibrosis and inflammation, reflected in a markedly reduced glomerular filtration rate (**Fig. 3a-e**).^28^ CIBERSORT analysis on RNAseq data over time indicated that lymphocytes were the most abundant immune cells is the kidney beyond the sixth month after injury while myeloid cell expansion characterized the acute phase (**Fig. 3f**).^29,30^ Beyond the sixth month after injury, lymphocytes were mostly organized in large cellular clusters, primarily around small arteries, between renal tubules identified by Havcr1 and Krt20 activity, markers of unresolved tubular injury (**Fig. 3g**, **Suppl. Fig. 1**).^28^ These highly vascularized ectopic lymphoid structures were populated by CD19^+^ B and CD3^+^ T cells (mainly CD4^+^) and surrounded by Lyve1^+^ lymphatic vessels.

**Figure 3.**
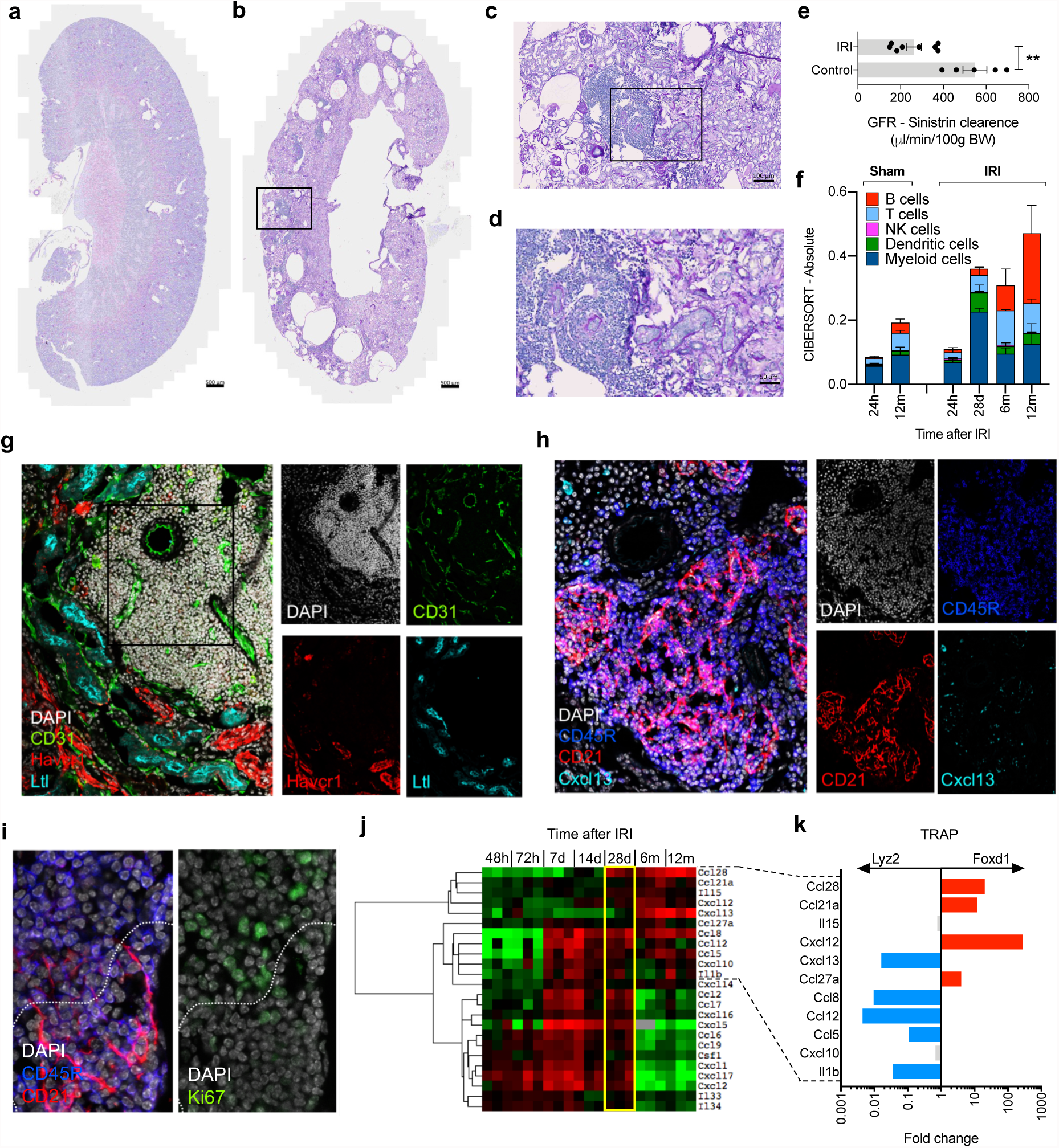
Tertiary lymphoid organs in the mouse kidney after ischemia/reperfusion injury. (a-d) Periodic acid-Schiff (PAS) staining of representative mouse kidney sections 16 months after IRI (b-d) or age-matched controls (a); n=3/group;scale bar=500 *µ*m in (a-b), 100 *µ*m in (c) and 50 *µ*m in (d). (e) Glomerular filtration rate (GFR), as measured by sinistrin clearence, in 7 mice 16-18 months after IRI and in 5 age-matched controls. **P<0.01. (f) Semi-quantitative evaluation of immune cell infiltrates in the kidney at different time points after IRI or sham surgery as determined by CIBERSORT analysis on whole kidney RNAseq data (n=3/group, *P<0.05). (g-h) Immunostaining of consecutive sections obtained from a representative mouse kidney 6 months after IRI (n=3-4/group). CD31: endothelial cells; Havcr1 (Kim1): marker of tubular injury; Ltl: proximal tubule. CD45R: B cells (and a subset of T cells, s. Suppl. Fig. 2); CD21 and Cxcl13: follicular dendritic cells. (i) Higher magnification of the B cell zone highlighting the partial separation of germinal centers in two areas:upper part with Ki67^+^ proliferating cells, lower part with CD21^+^ follicular dendritic cells. (j) Cluster analysis of cytokine transcripts obtained from RNAseq data from renal tissue at different time points after IRI (n=3 for each time point). (k) Cell specific expression analysis of cytokine transcripts from RNAseq data obtained after TRAP from Foxd1-derived renal stroma cells and Lyz2-derived myeloid cells at 28 days after IRI. The ratio of the mean RPKM values (Foxd1/Lyz2) obtained from 3 independent mice, including cytokine genes involved in the late immune response according to panel (j) are shown.Genes specifically expressed in myeloid cells are shown in blue and in stroma cells in red.

Examining the cellular compartmentalization of larger cellular aggregates identified a B cell zone with germinal centers of CD19+/CD45R+ or dim B cells embedded in a network of CD21^+^/Cxcl13^+^ follicular dendritic cells partially separated from clusters of highly proliferating Ki67^+^ lymphocytes, as typically observed in mature germinal centers (**Fig. 3h-i**).^31,32^ Similar ectopic lymphoid structures were described in different clinical conditions in the kidney^33^ and often develop in target tissues of autoimmune disease through the interaction of infiltrating immune cells and local stroma, directed by chemokine feedback loops.^34,35^ A systematic analysis of the cytokines transcribed in the injured kidney over time highlighted the progression from acute to chronic inflammation, with the late detection of cytokines involved in the formation of ectopic lymphoid tissue (e.g. Ccl21, Cxcl12, Cxcl13; **Fig. 3j**).^34^

At 28 days post IRI, the acute and the chronic inflammatory responses overlapped. To assess the contribution of different cellular compartments of the kidney at this critical transition point we performed RNA sequencing after Translating Ribosomal Affinity Purification (TRAP) to examine the profile of myeloid and renal stromal cell compartments (**Fig. 3k**).^36^ Both Ref34myeloid and stroma cells showed marked translation of cytokine encoding mRNAs. However, the stroma cell compartment, which encompasses the bulk of cells directly contributing to renal fibrosis,^37^ expressed a cluster of cytokines showing elevated expression sixth months after IRI, including cytokines associated with lymphocyte homing (e.g. Ccl28, Ccl21a and Cxcl12, **Fig. 3j-k**). Thus, adaptive immunity is an intrinsic component of dysfunctional kidney repair in the mouse kidney transitioning from acute to chronic injury in the absence of foreign antigens.

### B lymphocyte accumulation in the mouse kidney transitioning to chronic injury

The immune response detected in the kidney beyond the sixth month after IRI involved both T and B cells. The T cell compartment included a large fraction of non-conventional TCRαβ^+^CD4^-^CD8^-^ (double negative) T cells (**Suppl. Fig. 2**). Double negative T cells were predominant also among the lymphocytes isolated from the kidney of aged control mice, as shown in previous reports suggesting double negative T cells as a population of kidney resident T cells.^38,39^ Consistent with previous studies,^5,40^ B lymphocytes were rare in the normal kidney and memory B lymphocytes infiltrated the kidney in the first days after IRI; during the following weeks, B lymphocytes expanded massively and transcriptional analysis suggested a progressive switch to a plasma cell type (**Fig. 4a-b**). The direct phenotypic characterization of the infiltrating lymphocytes by flow cytometry after enzymatic tissue dissociation and magnetic cell sorting for CD45^42^ 16-18 months after IRI confirmed the presence of differentiated B lymphocytes, but showed that the renal B lymphocytes lacked CD138 (syndecan-1, a typical plasma cell marker, **Fig. 4c-d**, **Suppl. Fig. 3**). Instead of fully differentiated plasma cells, we consistently identified a population of CD19^+/low^CD45R^-^ B lymphocytes, that displayed CD126 (interleukin 6 receptor) and Cxcr4 (CD184) (**Fig. 4e-f, Suppl. Fig. 3**), the receptor for Cxcl12, a cytokine highly expressed by the renal stroma in the transition to chronic injury (**Fig. 3j-k**).

**Figure 4.**
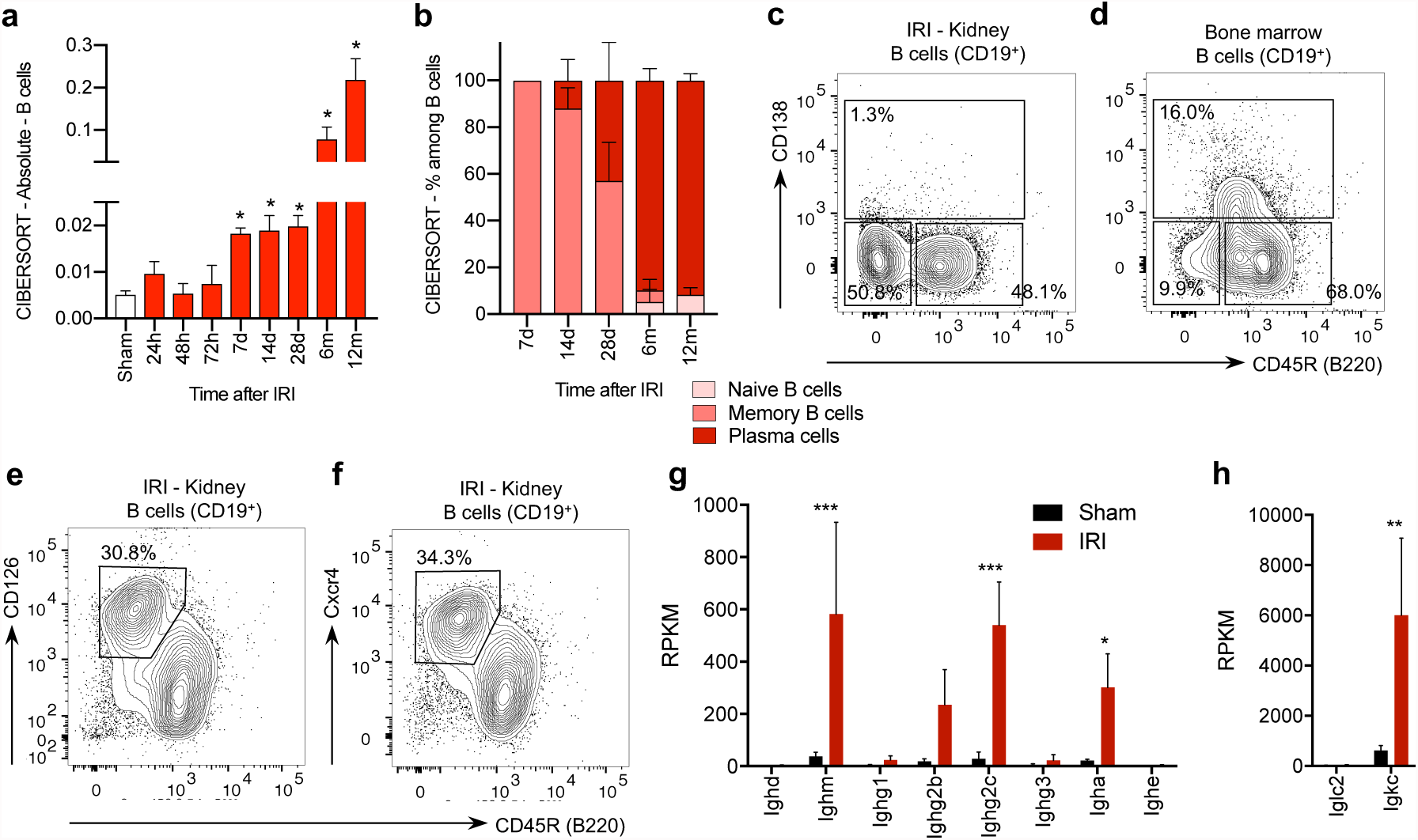
B lymphocytes in the late phase after ischemia-reperfusion injury in mice. (a-b) Semi-quantitative evaluation of immune cell infiltrates in the kidney at different time points after IRI or sham surgery as determined by CIBERSORT analysis on whole kidney RNAseq data (n=3/group, *P<0.05). (cf) Flow cytometric analysis on CD45+CD3-CD19+ ^or dim^ leukocytes isolated from the kidney or the bone marrow (BM) 16-18 months after IRI (1 representative example of 6 replicates is shown). (c-d) CD138 was absent on renal B cells, but detectable on the bone marrow B cells isolated from the same mouse (additional controls presented in Suppl. Fig. 3). (e-f) CD45+CD3-CD19+ ^or dim^CD45R^-^ B cells expressed high levels of CD126 and CD184 (Cxcr4). (g-h) RPKM values from RNAseq analysis on whole kidney 12 months after IRI or in sham controls focused on immunoglobulin constant regions transcripts (n=3/group, * P<0.05, *** P<0.001).

B lymphocytes isolated from the kidney in the late phase after IRI were reminiscent of plasma cell precursors involved in the renal production of autoantibodies in experimental lupus models and described in patients with systemic lupus erythematosus.^43^^-^^45^ The marked increase in immunoglobulin gene transcripts for IgM, IgG2c and IgA classes and Ig kappa light chains in whole organ RNA-profiles was consistent with the local production of antibodies in the kidney (**Fig. 4g-h**). Despite the high level of IgM transcripts and the recognized role of natural antibodies in the tissue injury response,^46^ B-1 cells (and marginal zone B cells) were not detected in the kidney (**Suppl. Fig. 4**). Thus, antibody secreting cells accumulated in the kidney in conjunction with the transition from acute to chronic kidney injury.

### B cell clonal expansion and affinity maturation in the injured mouse kidney

To better characterize the accumulation and the differentiation of B lymphocytes in the damaged mouse kidney we performed a B cell receptor (BCR) repertoire analysis in mice. First, we investigated the germline antibody gene segment repertoire by quantifying Ighv and Igkv transcripts in the kidney by whole organ RNA-seq. At 28 days after IRI we observed the relative enrichment of a limited number of Ighv genes, reflecting the presence of a small number of B cell. In the following months, in parallel with the massive increase of immunoglobulin transcripts, we detected the relative enrichment of V segments. The 3 most prevalent V segments accounted for 25-40% of the total Ighv transcripts and the top one for 18-35% of the Igkv transcripts at 12 months (**Fig. 5a-d**). This process could only be appreciated beyond the sixth month after IRI, underlining the need for a very long-term follow-up, and suggested the expansion of a subset of B cell clones in the kidney.

**Figure 5.**
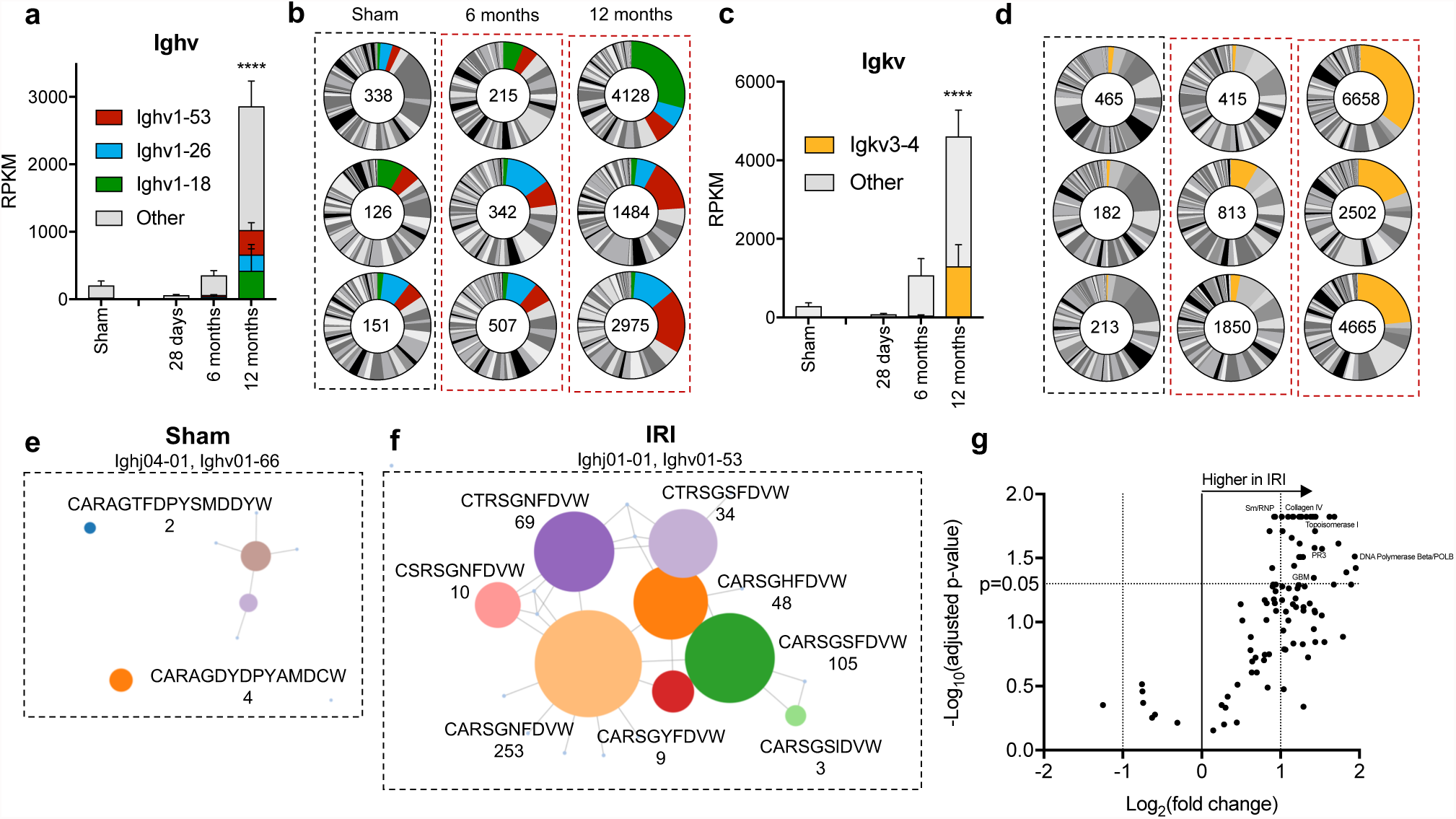
B cell receptor analysis and detection of autoantibodies. (a,c) Absolute RPKM values for germline Ighv and Igkv transcripts in 3 replicates collected 6 and 12 months after IRI and from sham control (matched to the 12 month IRI samples). The most frequent genes are highlighted with colors as indicated (n=3/group, ****P<0.0001 in comparison to sham control). (b,d) Relative frequency of germline Ighv and Igkv transcripts over time indicating the enrichment of common genes in the late phase; the number indicate the total RPKM of Ighv and Igkv at each time point. (e,f) Immunoseq^®^ analysis showing representatives cluster of vj rearrangements in sham and IRI mice (both 12 months after surgery).Each circle represent a unique amino acid sequence. The sequence is reported in the figure if >2 rearrangements were detected. The number of rearrangements is indicated below the amino acid sequence. The lines connect the circles if the corresponding rearrangements differ only in 1 amino acid. (g) Volcano plot comparison of signal to noise ratio in IgG reactivity in plasma samples collected 16-18 months after IRI (n=6) and in age-matched controls (n=6) on a microarray of autoantigens, with indication of log_2_-fold change on the x-axis (>0 indicates higher in IRI) and significance level, expressed as log_10_ of FDR-adjusted P-value. The name of representative antigens is indicated.

The most frequent segments in both Ighv and Ihkv were present at high frequency in the 3 mice analyzed at 12 months, but were not particularly abundant in sham control mice and were not among the most frequently used V segments in mice.^47^ This excludes an effect related to aging and suggest a biased usage of immunoglobulin genes. In a second step, the BCR repertoire analysis at 12 months after IRI was refined at the DNA level by Immunoseq^®^, which better assays the composition of the B cell repertoire providing additional information on the single VDJ rearrangements.^48^ This analysis confirmed the presence of a polyclonal B cell population in the kidney with the enrichment of a limited number of dominant clones. The amino acid sequences of the complementarity-determining region 3 (CDR3) of the immunoglobulin heavy chain in control mice did not show any evidence for a phylogenetic organization of the rearrangements (**Fig. 5e**). In contrast, IRI survivors presented clusters of hypermutated immunoglobulin gene rearrangements with few amino acid substitutions indicating a process of clonal expansion and affinity maturation (**Fig. 5f**). Thus, both histological findings and BCR analysis, provide evidence for the proliferation, selection and maturation of B lymphocytes in germinal centers within the kidney, in conjunction with the transition from acute to chronic kidney injury.

### Detection of autoantibodies after ischemia reperfusion injury

The clonal expansion of B lymphocytes in germinal centers is an antigen-triggered immunological process.^32^ In the absence of foreign antigens in this aseptic kidney injury model, we tested the plasma collected 16-18 months after IRI on a screening panel for autoantibodies. Thirty five out of 118 tested antigens showed a statistically significant increased signal to noise ratio (SNR) 16 months after IRI compared to controls (adjusted P-value<0.05, FDR 5%; **Fig. 5g**) without evidence for a common target antigen. This indicated that in the absence of foreign antigens, the intrarenal B cell response resulted in the production of broadly reactive autoantibodies. Since a similar process after kidney transplantation might stimulate the production of donor-specific antibodies, we retrospectively analyzed the clinical record of our study population to correlate the detection of anti-HLA antibodies with the initial injury response: among kidney recipients not HLA-sensitized at time point of transplantation. Indeed, 3 patients developed clinical evidence for late B cell mediated reactivity to the graft during further follow-up. All these patients were classified in the MIR group (**Suppl. Table 3**).

## Discussion

B lymphocytes are involved in the acute response to ischemic injury but are not essential for acute rejection, which is mainly mediated by T lymphocytes.^3-5,49^ In contrast, B cells are pivotal in the pathogenesis of late forms of immune-mediated graft injury. Here we show that late B cell activity in kidney allografts is tightly linked to dysfunctional kidney repair. Since in the clinical setting, this finding was associative, we turned to a mouse model to further characterize the immunological processes during the transition from acute to chronic kidney injury. A dysfunctional repair program following severe IRI transformed the mouse kidney into a tertiary lymphatic organ, which did not only host the inflammatory response typically associated to chronic organ injury,^50^ but also induced an antigen-driven expansion and maturation of B cell clones. In the absence of foreign antigens, this process resulted in the production of broadly reactive autoantibodies, as previously reported in patients after heart and brain injury.^51^^-^^53^

These findings are reminiscent of a recent characterization of the clonal evolution of autoreactive germinal centers in lupus. Once self-tolerance is broken in lupus, autoreactive B cells drive the expansion of further clones in the context of a self-sustained inflammatory feedback mechanism eventually leading to a convergent autoimmune response, with a delay of several months after the initial break of tolerance.^31^ Our findings support a model wherein dysfunctional tissue repair is sufficient to initiate a similar process with clinical relevance to organ transplantation.

Taken together, our data suggest a new model for the pathogenesis of late immune responses to allografts: a maladaptive repair program following unresolved tissue injury stimulates a chronic activation of the adaptive immune system, including B lymphocytes. B lymphocytes interact with the surrounding renal stroma to create an environment supporting the recruitment of additional clones and stimulating maturation and differentiation of antibody producing cells. Over time this process triggers the production of antibodies that promote further tissue damage in a deleterious feedback mechanism (**Fig. 6**).

**Figure 6.**
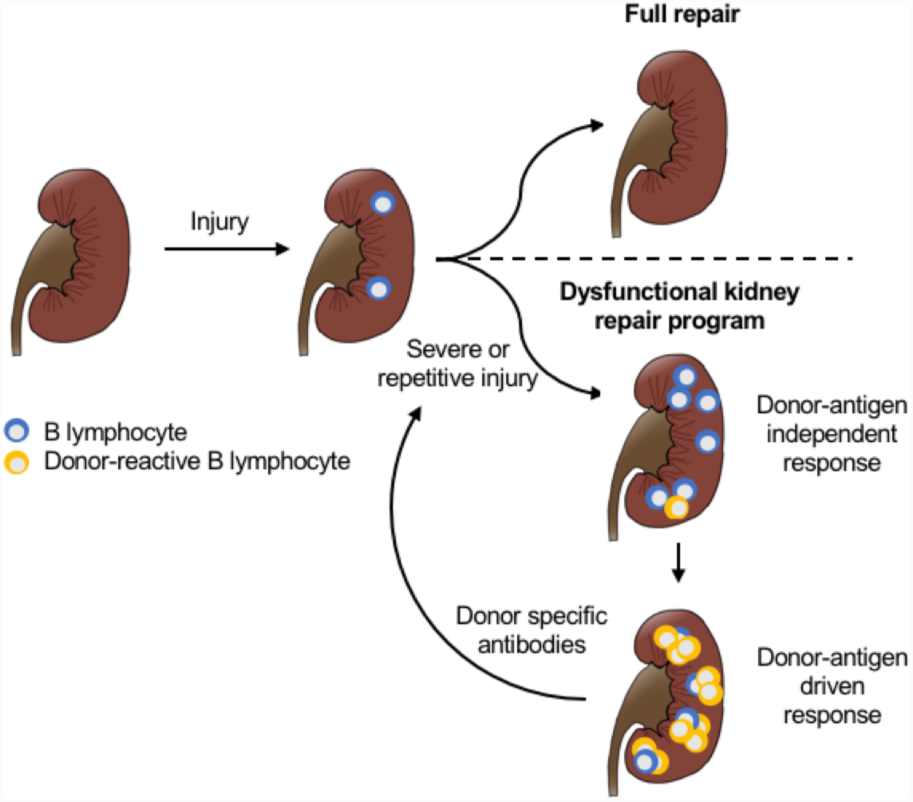
A new model to understand late immune responses to kidney allografts. Severe or repetitive kidney injury induces a dysfunctional repair program leading to a sustained immune response in the kidney.Over time, the local immune response leads to the recruitment and activation of donor reactive B cell clones, which differentiate to plasma cells and produce donor specific antibodies, further contributing to tissue injury in a deleterious feedback mechanism.

Although the numbers of patients was not sufficient to demonstrate a causal link to donor-specific immunoreactivity and to formally exclude potential confounders, the model is supported by earlier findings. Several studies characterized the broad antibody responses in conjunction with chronic rejection after kidney transplantation^54^ and the acceleration of chronic rejection by intragraft lymphoid neogenesis.^55^ Moreover, previous evidence for a stepwise breakdown of B cell tolerance as a preamble to chronic rejection directly parallels our observations in the mouse model.^56^ Also consistent with this model are reports showing that among the anti-HLA antibodies detected 5 years following transplantation, only one third were donor-specific, and non-donor specific antibodies appeared earlier than donor-specific, but both donor-specific antibodies and non-donor-specific antibodies were associated with an adverse outcome.^57,58^ Conversely, the identification of dysfunctional tissue repair as the initiator not only for fibrosis as previously proposed,^19^ but also for late forms of alloreactivity provides a plausible explanation for much lower incidence of chronic antibody-mediated rejection in an organ with a high regenerative capacity, such as the liver.^11^

Our model argues for a change in perspective in relation to the unresolved issue of chronic allograft dysfunction and B cell mediated immunity. In this, the production of donor-specific antibodies is a late event reflecting irreversible organ damage. Consequently, the lack of therapeutic efficacy against chronic antibody mediated rejection (including potent B cell depleting agents) is not surprising.^17^ However, our data – in line with previous clinical studies –^58^ indicate that the progression towards chronic damage and B cell activation is predictable, thereby offering an opportunity for early diagnostic and therapeutic intervention. The cause of injury in the clinical setting is much more variable than in the experimental model, several factors might determine if the repair response will result in a sustained immune activation, e.g. recent studies in mice suggest that age might play an important role in this regard.^59^ Developing strategies to attenuate the early injury response and repetitive hits during follow-up may prevent irreversible organ damage and late alloreactivity.^60^ The process characterized here may also have broader significance in the pathogenesis of chronic organ failure in other clinical conditions.^50,51^

## Acknowledgments

We thank Greg Alvarado, Jetty De Loor, Kari Koppitch, Gohar Seribekyan and Eric Tycksen for technical support. Work in APMs laboratory was supported by a grant from the California Institute for Regenerative Medicine (LA1-06536). PEC was supported by the Swiss National Science Foundation (grant 167773).

## Author contributions

PEC and APM contributed to conception and experimental design. JL performed IRI surgeries. PEC, JL, and SK performed experimental data acquisition and analysis. MN performed human data acquisition. PEC and BS performed human data analysis. PEC and APM prepared the manuscript, incorporating comments from other authors.

## Competing interests

None.

## Methods

### Studies in humans

#### Study design, patients and clinical data

We performed a genome-wide gene expression profile by RNA sequencing (RNAseq) in 42 kidney transplant recipients, randomly selected among patients with a full set of 4 available biopsies from a database at the University of Leuven. The patients received a kidney transplantation at the University of Leuven, Belgium. All patients gave written informed consent, and the study was approved by the Ethical Review Board of the University Hospital Leuven (S53364 and S59572). The renal biopsies were performed at the University Hospital of Leuven at following time points: before implantation (kidney flushed and stored on ice), 3 months and 12 months after transplantation (protocol biopsies). Additional biopsies performed in the same patients for a clinical indication were not considered for this study.

For histological evaluation, kidney sections were stained with hematoxylin eosin (HE), Periodic Acid-Schiff (PAS) and silver methenamine (Jones). All biopsies were centrally scored by pathologists dedicated to transplant pathology following the same standard procedures. The severity of chronic histological lesions was semi-quantitatively scored according to the Banff categories. The team involved in the computational analysis was not informed about any clinical information until the end of the study, when computational and clinical data were matched. The histological evaluation was independent of the computational analysis and the pathologist was not informed about the results of the transcriptional analysis.

#### Tissue storage and RNA extraction

Of each renal allograft biopsy included in this study, at least half a core was immediately stored on Allprotect Tissue Reagent^®^ (Qiagen Benelux BV, Venlo, The Netherlands), and after incubation at 4°C for at least 24 hours and maximum 72 hours, stored locally at -20°C, until shipment to the Laboratory of Nephrology of the KU Leuven. We performed RNA extraction using the Allprep DNA/RNA/miRNA Universal Kit^®^ (Qiagen Benelux BV, Venlo, The Netherlands) on a QIAcube instrument (Qiagen Benelux BV, Venlo, The Netherlands). The quantity (absorbance at 260nm) and purity (ratio of the absorbance at 230, 260 and 280nm) of the RNA isolated from the biopsies were measured using the NanoDrop ND-1000™ spectrophotometer (Thermo Scientific™, Life Technologies Europe BV, Ghent, Belgium). Before library preparation, RNA integrity was verified by high sensity RNA ScreenTape^®^ analysis. 5 samples were discarded because of not optimal RNA quality. Among the discarded samples, 3 were collected 12 months after transplantation, so that 39 patients could be classified in the MIR or non-MIR subgroups.

#### Library preparation, RNA sequencing and data processing

Library preparation and RNA sequencing were performed in two batches. The first batch consisted of a pilot study with 3 patients (12 samples) and the second batch included the rest of the study population. The library was prepared with Clontech SMARTer technology at the Genome Technology Access Center of the Washington University, St. Louis, MO. The sequencing was performed in the same lab by using the HiSeq 3000 system on the Illumina platform, with a target of 30M reads per sample. The reads were aligned to the Ensembl top-level assembly with STAR version 2.0.4b. Gene counts were derived from the number of uniquely aligned unambiguous reads by Subread:featureCount version 1.4.5. Transcript counts were produced by Sailfish version 0.6.3. Sequencing performance was assessed for total number of aligned reads, total number of uniquely aligned reads, genes and transcripts detected, ribosomal fraction known junction saturation and read distribution over known gene models with RSeQC version 2.3. All gene expression levels were normalized and quantified by RPKM (number of reads per kilobase per million mapped read).

### Studies in mice

#### Mice and surgical procedures

Mouse handling and husbandry and all surgical procedures were performed according to guidelines issued by the Institutional Animal Care and Use Committees (IACUC) at University of Southern California. Warm ischemia-reperfusion injury (IRI) was performed to induce ischemic acute kidney injury, as previously reported.^1^ Briefly, 10- to 12-week-old, 25- 28g, C57BL/6CN male mice, purchased from Charles River, were anesthetized with an intraperitoneal injection of a ketamine/xylazine (105 mg ketamine/kg; 10 mg xylazine/kg). Body temperature was maintained at 36.5-37°C throughout the procedure. The kidneys were exposed by a midline abdominal incision and both renal pedicles were clamped for 21 min using non-traumatic microaneurysm clips (Roboz Surgical Instrument Co.). Restoration of blood flow was monitored by the return of normal color after removal of the clamps. All the mice received intraperitoneal (i.p) 1 ml of normal saline at the end of the procedure. Sham-operated mice underwent the same procedure except for clamping of the pedicles.

#### Glomerular filtration rate

Glomerular filtration rate (GFR) was measured in mice by transcutaneous measurement of FITC-sinistrin disappearance with the NIC-Kidney device, purchased from Mannheim Pharma & Diagnostics GmbH (Mannheim, Germany), as previously reported. ^2^^-^^4^ Briefly, mice were anesthetized with isoflurane. The back was shaved and carefully depilated with a depilation cream to optimize the contact of the optical components of the detector with the skin. The depilation cream was carefully removed and the detector was placed directly on the skin and fixed with a tape. With the mouse still anesthetized, FITC-sinistrin (ca. 7.5 mg/100 g body weight, purchased from Mannheim Pharma & Diagnostics GmbH) was injected into the tail vein. After the injection the mouse was returned to its cage and transcutaneous detection of FITC-sinistrin disappearance was measured for 45 minutes. The data were analyzed with the NIC-Kidney device related software, which automatically provide the FITC-sinistrin half-life. GFR was calculated as follows:

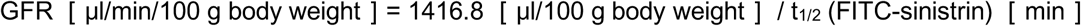

#### Histology and immunofluorescence

For conventional staining kidneys were perfused with ice-cold PBS and embedded in paraffin after overnight fixation in 4% paraformaldehyde (PFA) at 4°C. Sections were cut at 2 µm and stained with hematoxylin and eosin, or Periodic Acid-Schiff (PAS) staining at the histology core of the USC department of pathology, Los Angeles, CA. For immunofluorescence PFA fixed tissues were equilibrated in 30% sucrose/PBS overnight then embedded in OCT in dry ice ethanol bath. 8-10 µm frozen sections were washed in PBT (PBS + 1% Triton-X), blocked in in 5% Normal Donkey Serum in PBT, and incubated overnight at 4°C with primary antibodies and detected with species-specific secondary antibodies coupled to Alexa Fluor 488, 555, 594 and 647 (Life Technologies) for 1 hour at room temperature. Following antibodies were used in this study to recognize CD3 (rabbit, Abcam, ab16669), CD19 (rat, Invitrogen, 13-0194-82), CD21 (rabbit, Abcam, ab75985), CD31 (rat, BD, 553370), CD45 (goat, BD, 550280), CD45R (rat, BD, 557390), Cxcl13 (goat, R&D, AF470), F4/80 (rat, eBioscience, 14-4801), Havcr1 (goat, R&D, AF1817), Ltl lectin-FITC conjugate (Vector Laboratories, FL-1321), Lyve1 (goat, R&D, AF2125), Ki67 (rabbit, Novocastra, Ki67p-CE). All sections were stained with Hoechst 33342 (Life Technologies) prior to mounting with Immu-Mount (Fisher). All images were acquired on Zeiss Axio Scan Z1 slide scanner and Zeiss LSM780.

#### Cell isolation, sorting and flow cytometry

Lymphocyte were isolated from the kidney as previously described.^5^ Briefly, the mouse was perfused with PBS until the kidney was visually blood-free. The kidney was removed, mechanically dissociated and then incubated at 37°C in digestion buffer (RPMI medium, 1 mg/ml type I collagenase, 10 µg/ml DNAse, 10% FCS, 25 mM HEPES, penicillin and streptomycin). After 60 minutes the tissue was filtered through a 100 µm nylon mash to remove remaining large fragments. CD45^+^ cells were positively sorted by magnetic cell separation in an autoMACS with anti-mouse CD45 beads (Miltenyi biotec, Cat# 130-053-301). CD45^+^ cells were further characterized by flow cytometry (Aria II, BD bioscience). Following antibodies were purchased from BD bioscience: BV421-1B1 (CD1d), FITC-17A2 (CD3), APC and PE-RM4-5 (CD4), BV786-53-7.3 (CD5), APC53-6.7 (CD8a), PE-M1/70 (CD11b), BV605-HL3 (CD11c), FITC-1D3 (CD19), FITC and PE-1D3 (CD19), BV650-7G6 (CD21), APC-S7 (CD43), BV786-IM7 (CD44), APC-Cy7-30-F11 (CD45), APC-H7-RA3-6B2 (CD45R/B220), PE-DX5 (CD49b), PE-MEL-14 (CD62L), BV421-H1.2F3 (CD69), PE-D7715A7 (CD126), APC and BV786-281.2 (CD138), BV421-2B11/CXCR4 (CD184), BV421-Jo2 (Fas), APC and BV605-11-26c.2a (IgD), BV605-11/41 (IgM), BV421-R2-40 (IgG2a/b), BV786 and BV650-PK136 (NK1.1), PE-J43 (PD1), BV421-H5-597 (TCRβ), BV786-GL3 (TCRγδ). Cell viability was assessed by fixable viability staining 510.

#### RNA sequencing and Translating Ribosomal Affinity Purification

RNA was extracted from whole renal tissue or after Translating Ribosomal Affinity Purification, as previously reported,^1^ with a RNeasy^®^ kit (Qiagen) and provided to the USC Epigenome Center’s Data Production Core Facility for library construction and sequencing. RNA integrity was verified by Bio-Rad Experion analysis. Library construction was carried out using the Illumina TruSeq RNA Sample Prep kit v2 through polyA selection. The manufacturer’s protocol was followed with the exception that the final PCR amplification was performed for 12 and not 15 cycles. Libraries were visualized on the Agilent Bioanalzyer and quantified using the Kapa Biosystems Library Quantification Kit according to manufacturer’s instructions. Libraries were applied to an Illumina flow cell at a concentration of 16 pM on a version 3 flow cell and run on the Illumina HiSeq 2000 as a paired end read for 100 cycles each side. Image analysis and base calling was carried out using RTA 1.13.48.0. Final file formatting, de-multiplexing and fastq generation were carried out using CASAVA v 1.8.2. The sequencing data were aligned to mm10 genome assembly with STAR aligner (version 2.5.0b). The mapping index was generated using GENCODE release M4 (GRCm38.p3) gene annotations. In addition to read mapping, STAR was also used to remove duplicates, generate read-count tables and wiggle files. Differentially-expressed genes were called for each time point with DESeq2. Cluster analysis of cytokine transcripts was performed with Gene Cluster (version 3.0) and visualized with Treeview (version 1.1.6r4).

#### Autoantibodies

Autoantibody reactivity against a panel of autoantigens was determined in mouse plasma samples at the UT Southwestern Medical Center. The samples were incubated with autoantigen array and the autoantibodies binding to the antigens on the array was be detected with Cy3 labeled anti-IgG and Cy5 labeled anti-IgM to generate Tiff images. Genepix Pro 6.0 software was used to analyze the image. Signal-to-noise ratio (SNR) was used as a quantitative measure of the ability to resolve true signal from background noise. A higher SNR indicates higher signal over background noise. SNR equal or bigger than 3 was considered true signal from background noise. Quality control and normalization using variance stabilizing normalization were performed according to the standard procedure at UT Southwestern Medical center.

### General aspects

#### Statistical analyses

RPKM and eGFR values were compared by t-test or by Mann-Whitney U tests, a two-tailed P value <0.05 was considered significant. Categorical analyses were performed with Fisher’s exact or Chi-square tests. Differential gene expression analyses were performed with EdgeR,^6^ by applying an exact test, with a false discovery rate of 0.05. P values were adjusted according to the Benjamini-Hochberg procedures. SNR values from the autoantibodies analysis were compared by the two-stage step-up method of Benjamini, Krieger and Yekutiele with a false discovery rate of 0.05. Gene enrichment analyses were performed with ToppFun and P values were adjusted according to the Benjamini-Hochberg (https://toppgene.cchmc.org).

#### CIBERSORT and Immunoseq

CIBERSORT analyses were performed on RNAseq data by using the analytical tool developed by Newman and al. (cibersort.stanford.edu).^7^ LM22 was used as a reference signature for analysis on human data. The mouse specific reference profile developed by Chen et al. was applied for studies in mice.^8^

For Immunoseq^®^ analysis mouse renal tissue was analyzed by Adaptive biotechnologies (Seattle, USA).^9^ The data were analyzed with the immunoSEQ Analyzer (https://www.adaptivebiotech.com/immunoseq).

#### Software

Gene correlation analysis and feature correlation heatmaps were performed on the PIVOT platform, developed by the Kim Lab (http://kim.bio.upenn.edu/software/pivot.shtml).^10^ Dimension reduction analysis was performed with t-distributed stochastic neighbor embedding (t-SNE) by Rtsne package in R. Sample correlation analyses were performed on PIVOT^10^ or on Cluster (version 3.0) and visualized in TreeView (http://jtreeview.sourceforge.net). Box plots, scatter plots and histograms were generated with Prism 7.

---

**Suppl. figure 1.**
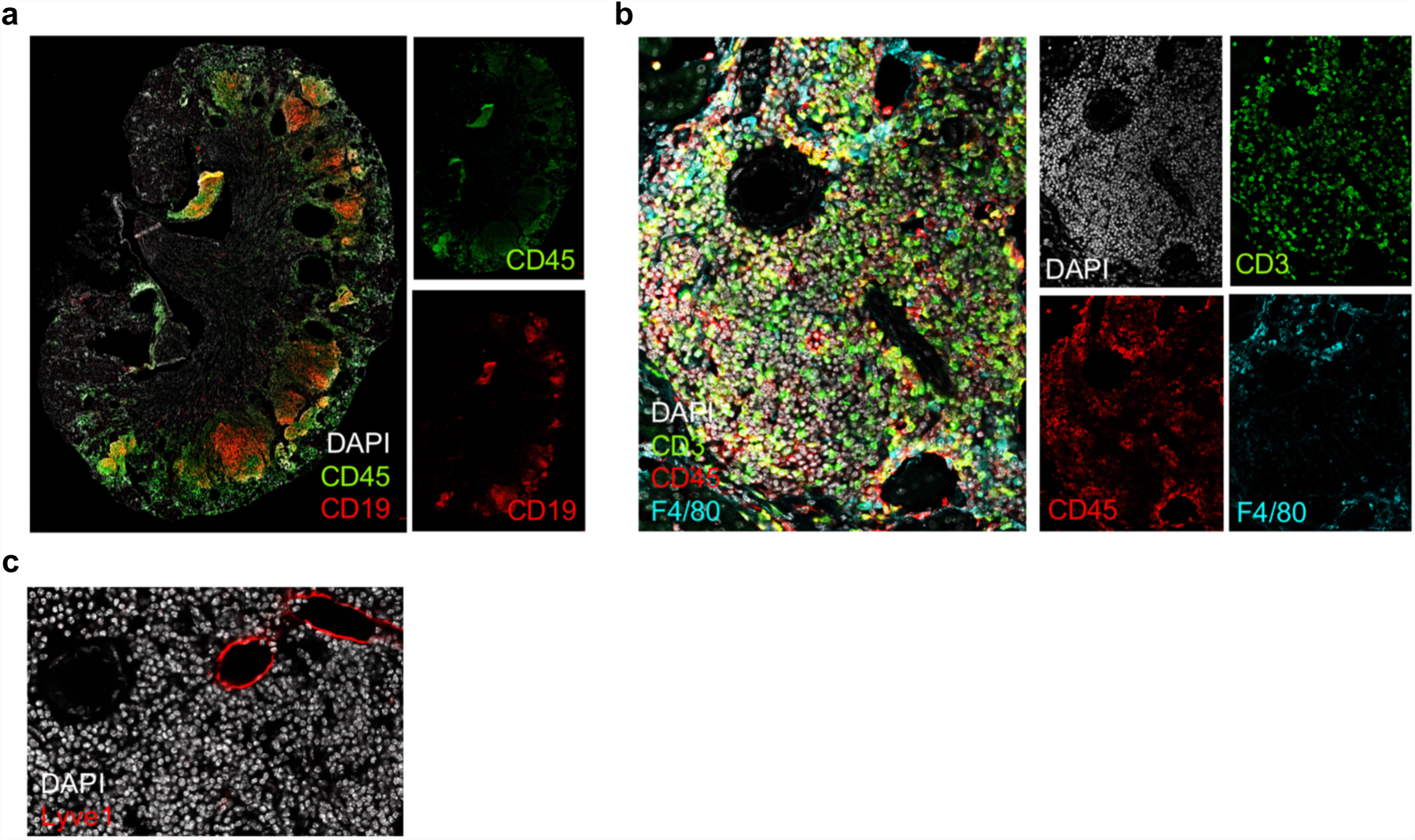
Characterization of renal ectopic lymphoid structures. Immunostaining of consecutive sections obtained from a representative mouse kidney 6 months after IRI (n=3-4/group). CD19: B cells; CD45: immune cells; CD3: T cells; F4/80: macrophages. CD45R: B cells (and a subset of T cells, s. suppl.figure 2); Lyve1: lymphatic vessels.

**Suppl. figure 2.**
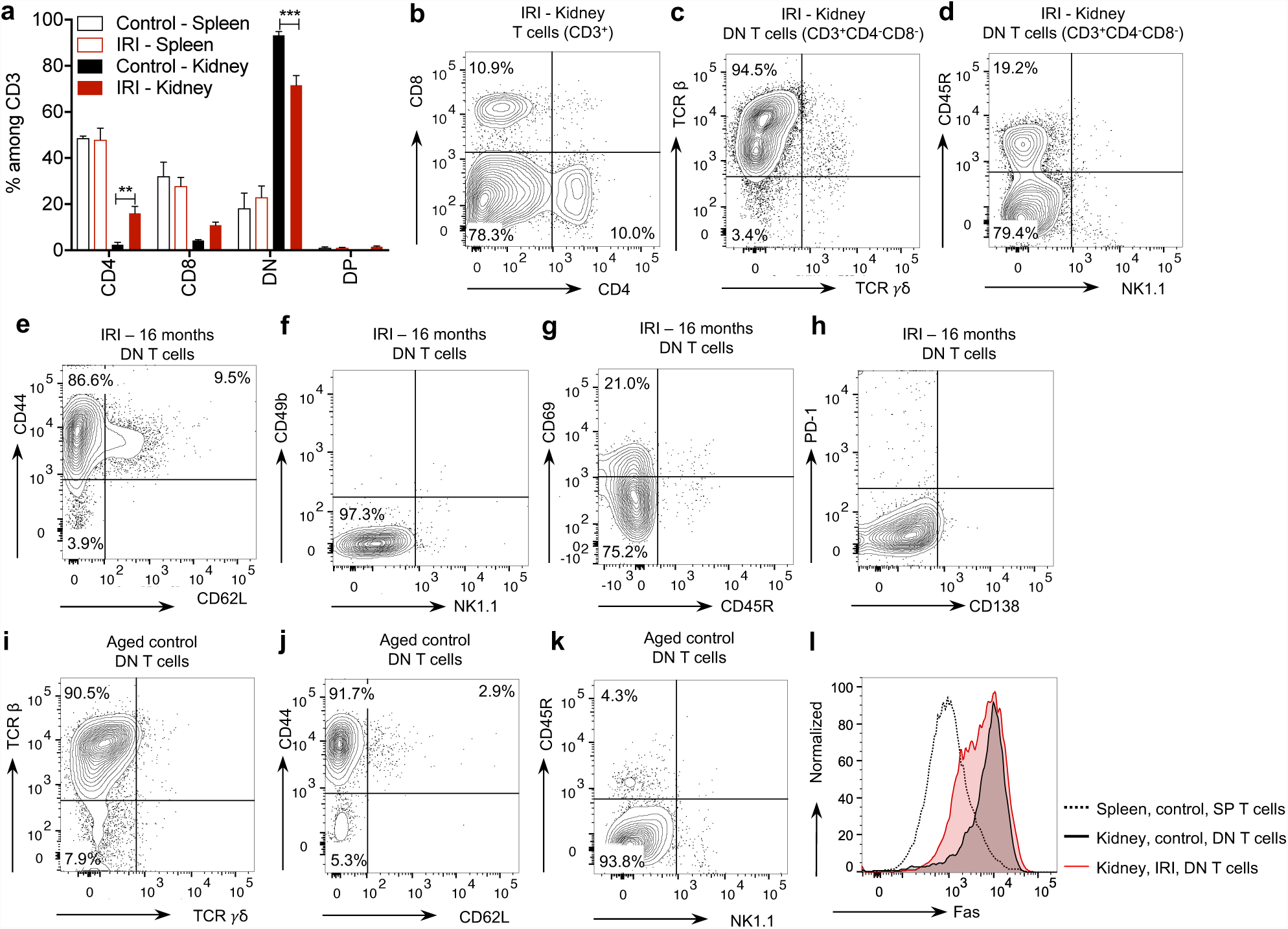
Characterization ofrenal Tcells after IRI. Flow cytometric analysis of leukocytes isolated from the kidney 16-18 months after IRI or from aged control mice, as indicated (representative examples are shown, n=6 in IRI and n=3 in controls). (a-b) General characterization of CD3^+^ T cells according to the expression of CD4 and CD8. DN: double negative, DP: double positive, ** P<0.01, *** P<0.001. (c-h) Further characterization of DN T cells. TCR: T cell receptor; NK1.1 and CD49b: natural killer cell-markers; CD45R is the isoform of CD45 typically expressed by B cells, and variably detectable on DN T cells. (i-k) Analogous characterization of renal DN T cells in age-matched control mice, showing a similar phenotype as detected after IRI, but not CD45R expression. (l) Representative flow cytometry analysis of Fas expression on T cells: DN T cells isolated from the kidney of both IRI and control mice, but not single positive (CD4 or CD8 positive) T cells from the spleen expressed high levels of the apoptosis receptor Fas.

**Suppl. figure 3.**
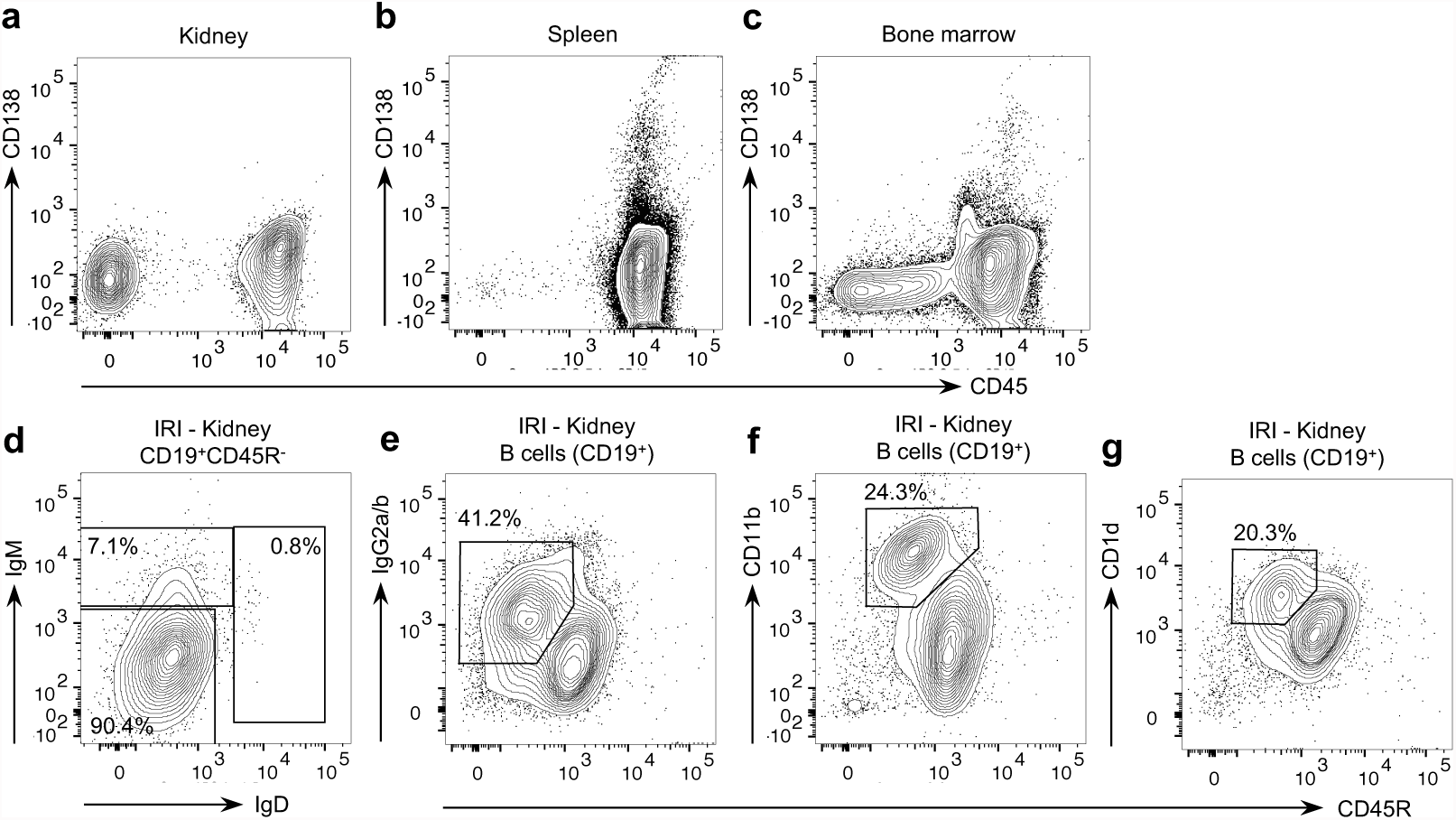
Characterization of renal B lymphocytes after IRI. Representative examples of flow cytometric analysis on leukocytes isolated from the kidney, the spleen and the bone marrow 16-18 months after IRI. (a-c) Controls to exclude that the absence of CD138 detection in renal B cells is related to the sorting procedure (compare with Fig. 4). Unsorted leukocytes isolated from the kidney, the spleen and the bone marrow and were directly stained for CD45 and CD138 without any additional manipulation. CD138 is detectable in the spleen and in the BM but not in the kidney. (d-g) Characterization of CD45^+^CD19^+^ B cells isolated from the kidney. (d) Only CD45R^- or dim^ cells are shown. (e-g) CD45R^- or dim^ cells are compared to CD45R^+^ cells. CD11b was reported in subsets of CD45R^-^ memory B cells (Driver et al., *J Immunol* 2001); and B cell expression of CD1d is involved in the pathogenesis of autoimmunity (Chaudhry et al., *J Immunol* 2014).

**Suppl. figure 4.**
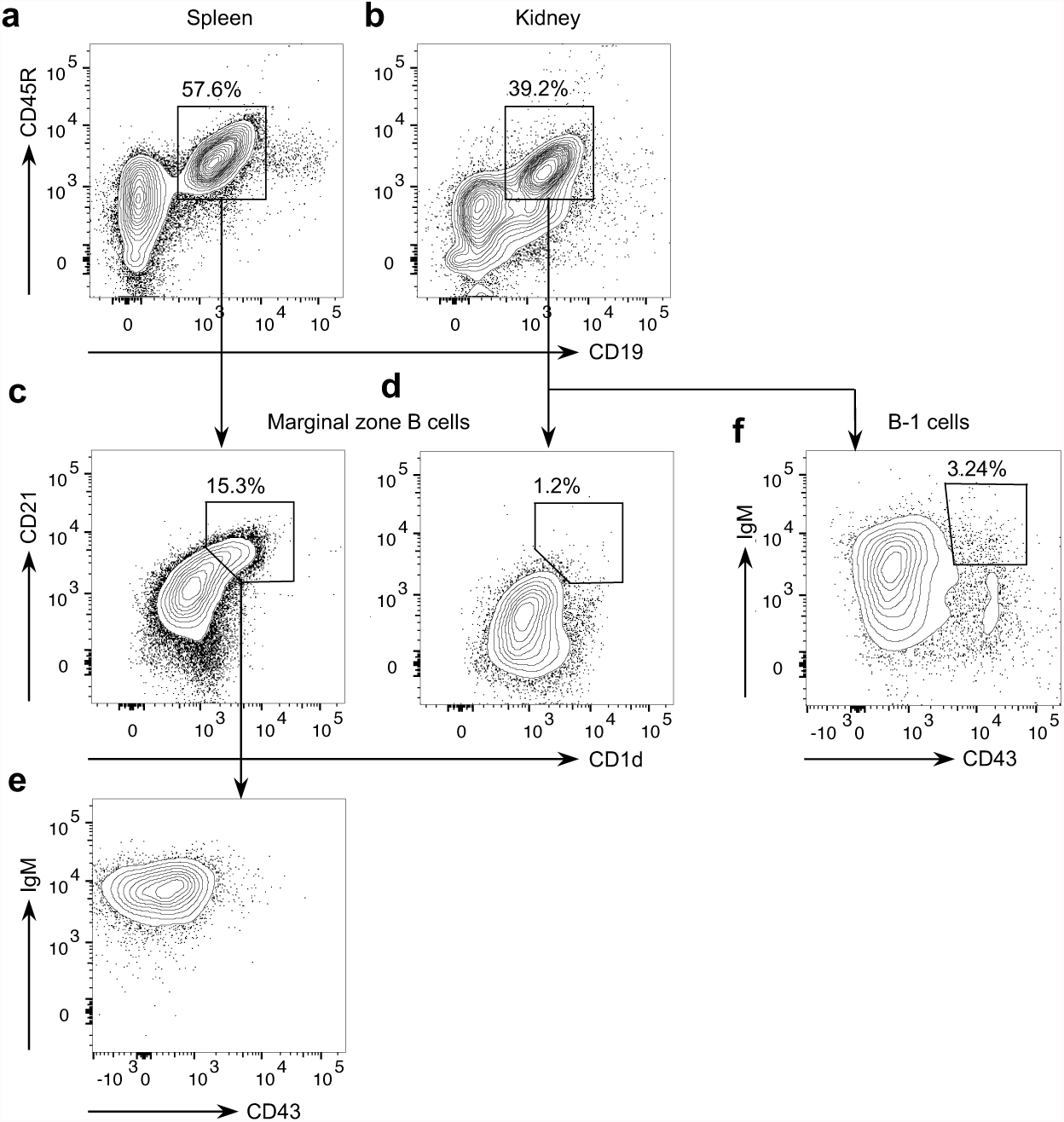
Flow cytometric analysis of leukocytes isolated from the kidney to investigate the presence marginal zone or B-1 cells after IRI. Cells isolated from the kidney (b,d,f) are compared with splenocytes (a,c,e). (c-e) Among CD45^+^CD19^+^CD45R^+^ isolated from the kidney we did not detect CD1d^+^CD21^+^ marginal zone B cells. In the spleen this B cell population was present and displayed the additional markers IgM and CD43. (f) CD43^+^IgM^+^ B-1 cells were not present in the kidney.

## References

1. Eming, S.A., Wynn, T.A. & Martin, P. Inflammation and metabolism in tissue repair and regeneration. Science 356, 1026-1030 (2017).

2. Jang, H.R. & Rabb, H. Immune cells in experimental acute kidney injury. Nat Rev Nephrol 11, 88-101 (2015).

3. Zouggari, Y., et al. B lymphocytes trigger monocyte mobilization and impair heart function after acute myocardial infarction. Nat Med 19, 1273-1280 (2013).

4. Chen, J., Crispin, J.C., Tedder, T.F., Dalle Lucca, J. & Tsokos, G.C. B cells contribute to ischemia/reperfusion-mediated tissue injury. J Autoimmun 32, 195-200 (2009).

5. Jang, H.R., et al. B cells limit repair after ischemic acute kidney injury. J Am Soc Nephrol 21, 654-665 (2010).

6. McDonald-Hyman, C., Turka, L.A. & Blazar, B.R. Advances and challenges in immunotherapy for solid organ and hematopoietic stem cell transplantation. Sci Transl Med 7, 280rv282 (2015).

7. Dai, H., et al. Donor SIRPalpha polymorphism modulates the innate immune response to allogeneic grafts. Sci Immunol 2(2017).

8. Celli, S., Albert, M.L. & Bousso, P. Visualizing the innate and adaptive immune responses underlying allograft rejection by two-photon microscopy. Nat Med 17, 744-749 (2011).

9. Sayegh, M.H. & Carpenter, C.B. Transplantation 50 years later--progress, challenges, and promises. N Engl J Med 351, 2761-2766 (2004).

10. Eskandary, F., et al. A Randomized Trial of Bortezomib in Late Antibody-Mediated Kidney Transplant Rejection. J Am Soc Nephrol (2017).

11. Valenzuela, N.M. & Reed, E.F. Antibody-mediated rejection across solid organ transplants: manifestations, mechanisms, and therapies. J Clin Invest 127, 2492-2504 (2017).

12. Wekerle, T., Segev, D., Lechler, R. & Oberbauer, R. Strategies for long-term preservation of kidney graft function. Lancet 389, 2152-2162 (2017).

13. Hariharan, S., et al. Improved graft survival after renal transplantation in the United States, 1988 to 1996. N Engl J Med 342, 605-612 (2000).

14. Meier-Kriesche, H.U., Schold, J.D. & Kaplan, B. Long-term renal allograft survival: have we made significant progress or is it time to rethink our analytic and therapeutic strategies? Am J Transplant 4, 1289-1295 (2004).

15. Colvin, R. B. & Smith, R.N. Antibody-mediated organ-allograft rejection. Nat Rev Immunol 5, 807-817 (2005).

16. Chong, A.S. & Ansari, M.J. Heterogeneity of memory B cells. Am J Transplant 18, 779-784 (2018).

17. Budde, K. & Durr, M. Any Progress in the Treatment of Antibody-Mediated Rejection? J Am Soc Nephrol 29, 350-352 (2018).

18. Einecke, G., et al. Expression of B cell and immunoglobulin transcripts is a feature of inflammation in late allografts. Am J Transplant 8, 1434-1443 (2008).

19. Venner, J.M., Famulski, K.S., Reeve, J., Chang, J. & Halloran, P.F. Relationships among injury, fibrosis, and time in human kidney transplants. JCI Insight 1, e85323 (2016).

20. Ferenbach, D.A. & Bonventre, J.V. Mechanisms of maladaptive repair after AKI leading to accelerated kidney ageing and CKD. Nat Rev Nephrol 11, 264-276 (2015).

21. Lagares, D., et al. ADAM10-mediated ephrin-B2 shedding promotes myofibroblast activation and organ fibrosis. Nat Med 23, 1405-1415 (2017).

22. Loupy, A., et al. The Banff 2015 Kidney Meeting Report: Current Challenges in Rejection Classification and Prospects for Adopting Molecular Pathology. Am J Transplant 17, 28-41 (2017).

23. Chiba, T., et al. Retinoic Acid Signaling Coordinates Macrophage-Dependent Injury and Repair after AKI. J Am Soc Nephrol 27, 495-508 (2016).

24. Kumar, S., et al. Sox9 Activation Highlights a Cellular Pathway of Renal Repair in the Acutely Injured Mammalian Kidney. Cell Rep 12, 1325-1338 (2015).

25. Modena, B. D., et al. Gene Expression in Biopsies of Acute Rejection and Interstitial Fibrosis/Tubular Atrophy Reveals Highly Shared Mechanisms That Correlate With Worse Long-Term Outcomes. Am J Transplant 16, 1982-1998 (2016).

26. Famulski, K.S., et al. Molecular phenotypes of acute kidney injury in kidney transplants. J Am Soc Nephrol 23, 948-958 (2012).

27. Cornell, L.D., Smith, R.N. & Colvin, R.B. Kidney transplantation: mechanisms of rejection and acceptance. Annu Rev Pathol 3, 189-220 (2008).

28. Liu, J., et al. Molecular characterization of the transition from acute to chronic kidney injury following ischemia/reperfusion. JCI Insight 2(2017).

29. Chen, Z., et al. Inference of immune cell composition on the expression profiles of mouse tissue. Sci Rep 7, 40508 (2017).

30. Newman, A.M., et al. Robust enumeration of cell subsets from tissue expression profiles. Nat Methods 12, 453-457 (2015).

31. Degn, S.E., et al. Clonal Evolution of Autoreactive Germinal Centers. Cell 170, 913-926 e919 (2017).

32. Victora, G.D., et al. Germinal center dynamics revealed by multiphoton microscopy with a photoactivatable fluorescent reporter. Cell 143, 592-605 (2010).

33. Segerer, S. & Schlondorff, D. B cells and tertiary lymphoid organs in renal inflammation. Kidney Int 73, 533-537 (2008).

34. Corsiero, E., Nerviani, A., Bombardieri, M. & Pitzalis, C. Ectopic Lymphoid Structures: Powerhouse of Autoimmunity. Front Immunol 7, 430 (2016).

35. Krautler, N.J., et al. Follicular dendritic cells emerge from ubiquitous perivascular precursors. Cell 150, 194-206 (2012).

36. Liu, J., et al. Cell-specific translational profiling in acute kidney injury. J Clin Invest 124, 1242-1254 (2014).

37. Grgic, I., et al. Translational profiles of medullary myofibroblasts during kidney fibrosis. J Am Soc Nephrol 25, 1979-1990 (2014).

38. Mayans, S., et al. alphabetaT cell receptors expressed by CD4(-)CD8alphabeta(-) intraepithelial T cells drive their fate into a unique lineage with unusual MHC reactivities. Immunity 41, 207-218 (2014).

39. Martina, M.N., et al. Double-Negative alphabeta T Cells Are Early Responders to AKI and Are Found in Human Kidney. J Am Soc Nephrol 27, 1113-1123 (2016).

40. Park, J., et al. Single-cell transcriptomics of the mouse kidney reveals potential cellular targets of kidney disease. Science (2018).

41. Shi, W., et al. Transcriptional profiling of mouse B cell terminal differentiation defines a signature for antibody-secreting plasma cells. Nat Immunol 16, 663-673 (2015).

42. Ludwig-Portugall, I. & Kurts, C. T cell isolation from mouse kidneys. Methods Mol Biol 1193, 27-35 (2014).

43. McHeyzer-Williams, L.J., Cool, M. & McHeyzer-Williams, M.G. Antigen-specific B cell memory: expression and replenishment of a novel b220(-) memory b cell compartment. J Exp Med 191, 1149-1166 (2000).

44. Arce, E., et al. Increased frequency of pre-germinal center B cells and plasma cell precursors in the blood of children with systemic lupus erythematosus. J Immunol 167, 2361-2369 (2001).

45. Espeli, M., et al. Local renal autoantibody production in lupus nephritis. J Am Soc Nephrol 22, 296-305 (2011).

46. Fleming, S.D. & Tsokos, G.C. Complement, natural antibodies, autoantibodies and tissue injury. Autoimmun Rev 5, 89-92 (2006).

47. Arnaout, R., et al. High-resolution description of antibody heavy-chain repertoires in humans. PLoS One 6, e22365 (2011).

48. Carlson, C.S., et al. Using synthetic templates to design an unbiased multiplex PCR assay. Nat Commun 4, 2680 (2013).

49. Rauch, P.J., et al. Innate response activator B cells protect against microbial sepsis. Science 335, 597-601 (2012).

50. Netea, M.G., et al. A guiding map for inflammation. Nat Immunol 18, 826-831 (2017).

51. Dick, S.A. & Epelman, S. Chronic Heart Failure and Inflammation: What Do We Really Know? Circ Res 119, 159-176 (2016).

52. Iadecola, C. & Anrather, J. The immunology of stroke: from mechanisms to translation. Nat Med 17, 796-808 (2011).

53. Li, X., et al. Autoantibody profiling on a plasmonic nano-gold chip for the early detection of hypertensive heart disease. Proc Natl Acad Sci U S A 114, 7089-7094 (2017).

54. Porcheray, F., et al. Chronic humoral rejection of human kidney allografts associates with broad autoantibody responses. Transplantation 89, 1239-1246 (2010).

55. Sicard, A., Chen, C.C., Morelon, E. & Thaunat, O. Alloimmune-induced intragraft lymphoid neogenesis promotes B-cell tolerance breakdown that accelerates chronic rejection. Curr Opin Organ Transplant 21, 368-374 (2016).

56. Thaunat, O., et al. A stepwise breakdown of B-cell tolerance occurs within renal allografts during chronic rejection. Kidney Int 81, 207-219 (2012).

57. Hourmant, M., et al. Frequency and clinical implications of development of donor-specific and nondonor-specific HLA antibodies after kidney transplantation. J Am Soc Nephrol 16, 2804-2812 (2005).

58. O’Connell, P.J., et al. Biopsy transcriptome expression profiling to identify kidney transplants at risk of chronic injury: a multicentre, prospective study. Lancet 388, 983-993 (2016).

59. Sato, Y., et al. Heterogeneous fibroblasts underlie age-dependent tertiary lymphoid tissues in the kidney. JCI Insight 1, e87680 (2016).

60. Nasralla, D., et al. A randomized trial of normothermic preservation in liver transplantation. Nature 557, 50-56 (2018).

61. Byun, J.S., Park, S., Caban, A., Jones, A. & Gardner, K. Linking Race, Cancer Outcomes, and Tissue Repair. Am J Pathol 188, 317-328 (2018).

## References

1. Liu, J, Krautzberger, AM, Sui, SH, Hofmann, OM, Chen, Y, Baetscher, M, Grgic, I, Kumar, S, Humphreys, BD, Hide, WA, McMahon, AP: Cell-specific translational profiling in acute kidney injury. J Clin Invest, 124: 1242-1254, 2014.

2. Schock-Kusch, D, Xie, Q, Shulhevich, Y, Hesser, J, Stsepankou, D, Sadick, M, Koenig, S, Hoecklin, F, Pill, J, Gretz, N: Transcutaneous assessment of renal function in conscious rats with a device for measuring FITC-sinistrin disappearance curves. Kidney Int, 79: 1254-1258, 2011.

3. Herrera Perez, Z, Weinfurter, S, Gretz, N: Transcutaneous Assessment of Renal Function in Conscious Rodents. J Vis Exp: e53767, 2016.

4. Schock-Kusch, D, Geraci, S, Ermeling, E, Shulhevich, Y, Sticht, C, Hesser, J, Stsepankou, D, Neudecker, S, Pill, J, Schmitt, R, Melk, A: Reliability of transcutaneous measurement of renal function in various strains of conscious mice. PLoS One, 8: e71519, 2013.

5. Ludwig-Portugall, I, Kurts, C: T cell isolation from mouse kidneys. Methods Mol Biol, 1193: 27-35, 2014.

6. Robinson, MD, Smyth, GK: Small-sample estimation of negative binomial dispersion, with applications to SAGE data. Biostatistics, 9: 321-332, 2008.

7. Newman, AM, Liu, CL, Green, MR, Gentles, AJ, Feng, W, Xu, Y, Hoang, CD, Diehn, M, Alizadeh, AA: Robust enumeration of cell subsets from tissue expression profiles. Nat Methods, 12: 453-457, 2015.

8. Chen, Z, Huang, A, Sun, J, Jiang, T, Qin, FX, Wu, A: Inference of immune cell composition on the expression profiles of mouse tissue. Sci Rep, 7: 40508, 2017.

9. Carlson, CS, Emerson, RO, Sherwood, AM, Desmarais, C, Chung, MW, Parsons, JM, Steen, MS, LaMadrid-Herrmannsfeldt, MA, Williamson, DW, Livingston, RJ, Wu, D, Wood, BL, Rieder, MJ, Robins, H: Using synthetic templates to design an unbiased multiplex PCR assay. Nat Commun, 4: 2680, 2013.

10. Zhu, Q, Fisher, SA, Dueck, H, Middleton, S, Khaladkar, M, Kim, J: PIVOT: platform for interactive analysis and visualization of transcriptomics data. BMC Bioinformatics, 19: 6, 2018.

